# The Effects of Prenatal Nicotine and Thc E-Cigarette Exposure on Motor Development in Rats

**DOI:** 10.1101/2021.10.20.465160

**Authors:** S. Hussain, K. R. Breit, J. D. Thomas

## Abstract

In the United States, nicotine and cannabis are the most common licit and illicit drugs used among pregnant women. Importantly, nicotine and cannabis are now being combined for consumption via e-cigarettes, an increasingly popular route of administration. Both nicotine and tetrahydrocannabinol (THC), the primary psychoactive component of cannabis, cross the placenta barrier. However, the consequences of prenatal cannabis use are not well understood, and less is known about potential combination effects when consumed with nicotine, especially via e-cigarettes. The present study used a rodent model to examine how prenatal e-cigarette exposure to nicotine, THC, and the combination impacts motor development among offspring. Pregnant Sprague-Dawley rats were exposed to nicotine (36 mg/mL), THC (100 mg/mL), the combination, or vehicle via e-cigarette inhalation from gestational days (GD) 5-20. One sex pair per litter was tested on an early sensorimotor development task (postnatal days [PD] 12-20) and a parallel bar motor coordination task (PD 30-32). Combined prenatal exposure to nicotine and THC delayed sensorimotor development, even though neither drug produced impairments on their own. In contrast, prenatal exposure to either nicotine or THC impaired motor coordination, whereas combined exposure exacerbated these effects, particularly among females. These data illustrate that prenatal exposure to either nicotine or THC may alter motor development, and that the combination may produce more severe effects. These findings have important implications for pregnant women as we better understand the teratogenic effects of these drugs consumed via e-cigarettes.

## INTRODUCTION

Prevalence rates of tobacco use during pregnancy are estimated to be around 16% (Forray, 2016). Both clinical and preclinical research have shown that cigarette smoking during pregnancy can disrupt fetal development, exerting long-lasting physical, neurodevelopmental, cognitive, and behavioral effects (Ernst et al., 2001; Holz et al., 2014; Tiesler & Heinrich, 2014). Tobacco and nicotine use during pregnancy has been associated with increased risk of miscarriage, sudden infant death syndrome, preterm birth, low birth weight, and impaired lung development (Ernst et al., 2001; Lambers & Clark, 1996; Taghavi et al., 2018; Wickstrom, 2007). Behaviorally, prenatal exposure to tobacco and/or nicotine has also been associated with poorer academic performance (Blood-Siegfried & Rende, 2010; Longo et al., 2014), impaired learning and memory, increased risk for attention deficits, internalizing and externalizing behaviors, substance abuse and other mental health disorders (Holbrook, 2016; Hunter et al., 2020; Tiesler & Heinrich, 2014). Consistent with the clinical literature, rodent studies have shown that prenatal nicotine exposure can reduce birth weight, disrupt CNS development, and lead to delays in reflex development, impaired learning and memory, increased anxiety, increased risk for later substance use, and other affective alterations (Ajarem & Ahmad, 1998; Blood-Siegfried & Rende, 2010; Ernst et al., 2001; Lee & Picciotto, 2019; Tizabi et al., 2000). Importantly, results vary depending on administration route, developmental timing, and dose (Polli & Kohlmeier, 2020).

However, nicotine is frequently not the only drug consumed during pregnancy; cannabis is the most commonly used illicit drug among pregnant women in the United States, with prevalence rates estimated to be between 5-10% (Brown et al., 2017; Thompson et al., 2019). Importantly, since cannabis use has become legalized in many states, cannabis products have become more available, and use continues to increase, even among pregnant women and women of reproductive age (Hall & Weier, 2015). For example, a recent study found 21% of women from a sample in Northern California used cannabis during their pregnancy (Young-Wolff et al., 2019). In addition to recreational use, women have reported using cannabis while pregnant to relieve nausea, as well as to reduce pain and anxiety (Thompson et al., 2019).

Compared to prenatal nicotine exposure, the consequences of prenatal cannabis exposure are much less understood. Although prenatal cannabis exposure generally does not induce robust physical effects, cannabis use during pregnancy is associated with decreased gestational length, reduced birth weight, and higher likelihood of admittance to the NICU (Amlani et al., 2017; Goldschmidt et al., 2016; Grant et al., 2018; Gunn et al., 2015; Paul et al., 2019). Clinical research suggests that prenatal cannabis use can induce long-term adverse effects on brain, cognitive, and behavioral development of the exposed child, although results have been mixed. Effects of prenatal cannabis exposure may be evident even in newborns, who exhibit changes in arousal (Dahl et al., 1995; Scher et al., 1988), and children exposed prenatally to cannabis exhibit poorer academic performance, with deficits in verbal skills, memory, and attention (Fried, 1991; Fried, 1996; Fried et al., 1998; Huizink, 2014; Smith et al., 2016). In addition, prenatal cannabis is associated with reduced impulse control, affective disorders, disrupted sleep (Nashed et al., 2020), as well as increased risk for psychopathology (Paul et al., 2019). As delta-9-tetrahydrocannabinnol (THC) potency has risen sharply in recent years (Chandra et al., 2019), the consequences of high levels of THC exposure may be more severe and remain largely unknown.

Although results have also been mixed among preclinical studies, accumulating evidence suggests that maternal exposure to various cannabinoid compounds can adversely affect physical, cognitive, affective, and social development. For example, prenatal cannabinoid exposure can disrupt eye development (Fish et al., 2017; Fish et al., 2016) and restrict body growth (Natale et al., 2020). Behaviorally, prenatal THC exposure can increase anxiety-like behavior, emotional reactivity, and alter social behavior (Newsom & Kelly, 2008; Weimar et al., 2020). Early exposure to THC or other cannabinoids also can lead to impairments in learning and memory (Riedel & Davies, 2005), motor development (Breit et al., 2019), as well as a host of other behavioral alterations (Campolongo et al., 2011).

In addition, 50% of women in the United States who report drug use during pregnancy engage in polydrug consumption, with the most common combination being nicotine and cannabis (Forray, 2016). In fact, numerous studies suggest that maternal co-use of nicotine and cannabis is on the rise (Coleman-Cowger et al., 2017). Increased rates of co-use of nicotine and THC has been partly due to use of electronic cigarettes (e-cigarettes) or vaping (Coleman-Cowger et al., 2018; Glasser et al., 2017), which allow for easy combinations of drugs. The United States has the largest and quickest growing market for e-cigarettes (McCubbin et al., 2017) and many consumers, including pregnant women (McCubbin et al., 2017), assume that e-cigarettes are a safer alternative than traditional combustible cigarettes (Glasser et al., 2017). However, there is limited research exploring whether e-cigarette use is safe during pregnancy, and how the combination of prenatal exposure to nicotine and THC may influence development.

Using a rat model, this study examined whether prenatal nicotine and/or THC exposure via e-cigarettes in pregnant rats affected motor development among offspring. Pregnant dams were exposed to nicotine, THC, the combination, or vehicle via vapor inhalation. Following birth, offspring of the exposed dams were tested on an early sensorimotor development task (postnatal days (PD) 12-20) and a motor coordination task later in early adolescence (PD 30-32).

## METHODS

All procedures and behavioral testing measures used in this experiment were approved by the Institutional Animal Care and Use (IACUC) committee at San Diego State University (SDSU) and are in accordance with the National Institute of Health Guide for the Care and Use of Laboratory Animals.

### Prenatal Vapor Exposure

Breeding and testing procedures took place at the Center for Behavioral Teratology at San Diego State University. The study design is depicted in Figure 1. One adult male and one adult female Sprague-Dawley rat were housed together overnight; the presence of a seminal plug suggested that mating occurred and was designated as gestational day (GD) 0. Pregnant dams were randomly assigned to receive exposure to nicotine (36 mg/mL; Sigma-Aldrich), THC (100 mg/mL; NIDA Drug Supply Program), the combination, or the e-cigarette vehicle (propylene glycol; Sigma-Aldrich) daily from GD 5-20 during a 40-min vapor inhalation session (2 L/min airflow; 1 6-sec puff every 5 min for 30 minutes followed by a 10-min post-exposure air clearance). The vapor inhalation equipment was designed by La Jolla Alcohol Research Inc and used commercially available e-cigarette tanks (SMOK V8 X-Baby Q2). This exposure did not significantly affect maternal weight gain, daily food and water intake, litter size, birth weight, or litter sex ratio (Breit et al., 2021).

**Fig. 1.**
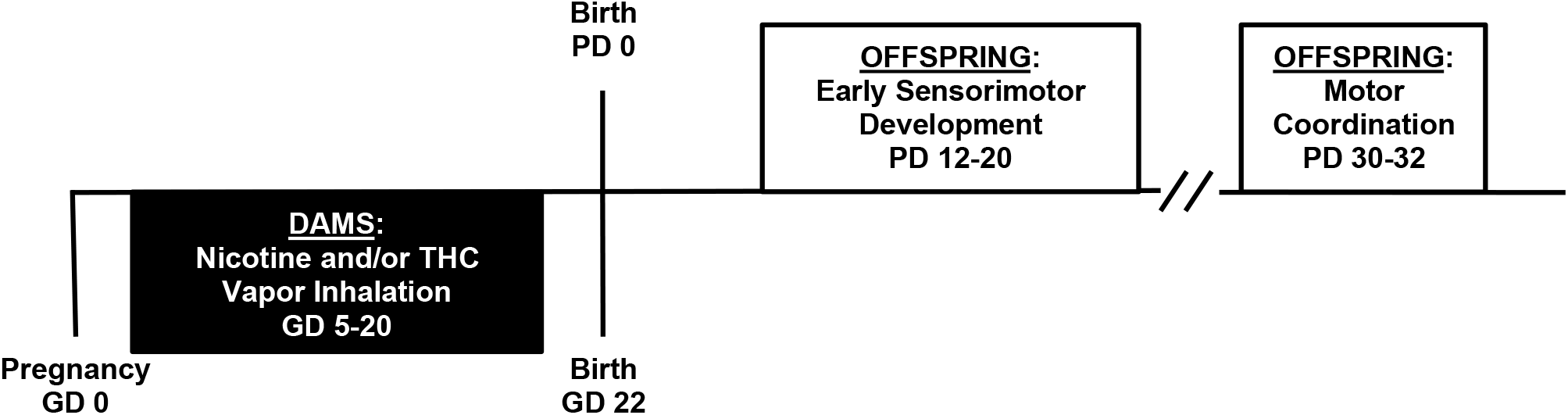
Timeline of study design

### Offspring Subjects

To control for possible litter effects, only one sex pair from each litter was pseudorandomly assigned to complete both motor tasks. Remaining offspring were used for additional behavioral testing that will be reported separately. Body weights were recorded on each day of behavioral testing, and subjects were acclimated to the testing rooms for 30 min before any testing began.

### Grip Strength and Hindlimb Coordination Task

A grip strength and hindlimb coordination task was used to measure early sensorimotor development from PD 12-20. Pups were suspended by their forepaws on a wire (2-mm diameter) above a cage of bedding for two consecutive trials each day. Trials were deemed successful if pups were able to either hold onto the wire for 30-sec or place their hindlimb on the wire. The first day of a successful trial, the percent of successful subjects each day, and the number of successful trials were recorded live by two investigators; all testing was videorecorded for any additional future analyses.

### Parallel Bar Motor Coordination Task

Motor coordination and balance was tested in subjects using a parallel bar task from PD 30-32. The parallel bar apparatus consists of two parallel steel rods (0.5 cm diameter, 91 cm long) held between 2 end platforms (15.5 × 17.8 cm), above a cage of bedding. The width between the bars could be expanded by 0.5-cm increments; the initial width was set at 4.0 cm. To minimize stress and promote movement, motor coordination testing was performed under a dim red light.

Subjects were placed on each platform to acclimate for 30 seconds; after acclimation, subjects were placed in the middle of the parallel rods, with left paws on one rod and right paws on the other, to observe traversals to the platform. A successful trial consisted of four successive alternating steps with their left and right hindlimb paws. Once a successful trial was completed, the width between rods was increased by 0.5 cm. If the subject fell off the rods, swung under the bars, dragged a paw, or otherwise failed to take four coordinated successive steps, the trial was deemed unsuccessful and the subject was placed back on the rods for another trial. If the subject had 5 unsuccessful trials in a row, they were returned to their home cage and testing was complete for that day. Subjects were allowed a maximum of 15 trials each day. On each subsequent day, testing was resumed with each subject beginning on the last successful width recorded for that subject. The number of trials to first success, maximum width achieved, and success ratio (successful trials divided by total trials attempted) were recorded each day.

### Data Analyses

All data were analyzed using SPSS software version 27 (SPSS Inc., Chicago, IL) with significance levels set at *p* < 0.05. For data that were normally distributed (Shapiro-Wilk test), data were initially analyzed using 2 (Nicotine) × 2 (THC) × 2 (Sex) ANOVAs. Additional ANOVAs using exposure group as a factor (Nicotine+THC, Nicotine, THC, Vehicle) were used with post hoc tests (Fisher’s Least Significant Difference), when appropriate. For data collected across days, repeated measure ANOVAs were used. For data that were not normally distributed or were binomial, non-parametric analyses were used.

## RESULTS

### Early Sensorimotor Development

A total of 96 subjects were tested on the early sensorimotor development task (Nicotine+THC [NIC+THC]: 12 females, 12 males; Nicotine [NIC]: 13 females, 11 males; THC [THC]: 12 females, 10 males; Vehicle [VEH]: 14 females, 12 males). Data from the grip strength and hindlimb coordination task were not normally distributed (Shapiro Wilk tests: all *p*’s < 0.05); thus, non-parametric analyses were used. Mann-Whitney analyses were used to determine main effects of prenatal nicotine exposure (Nicotine, Vehicle), prenatal THC exposure (THC, Vehicle), and sex (female, male). No differences were detected between male and female offspring, thus all data are shown collapsed by sex. To examine potential interactive effects of prenatal nicotine and THC exposure, Kruskal-Wallis analyses were also used to examine group differences (Nicotine+THC, Nicotine, THC, Vehicle), with follow-up chi-square analyses, when appropriate.

### Body Weights

Subjects exposed to THC alone tended to weigh more than the nicotine-exposed groups, producing a main effect of nicotine (F[1,88] = 4.08, *p* < 0.05) as well as a Day*Nicotine interaction (F[8,704] = 2.15, *p* < 0.05). Subjects exposed to prenatal nicotine weighed less than those not exposed to nicotine on PD 12, 13, 15, 16, 19, and 20; however, this effect was driven by nicotine-exposed subjects weighing less than subjects exposed to THC alone (all *p*’s < 0.05), and not controls (Figure 2). In fact, when comparing all four groups to one another, there were no significant differences in body weight.

**Fig. 2.**
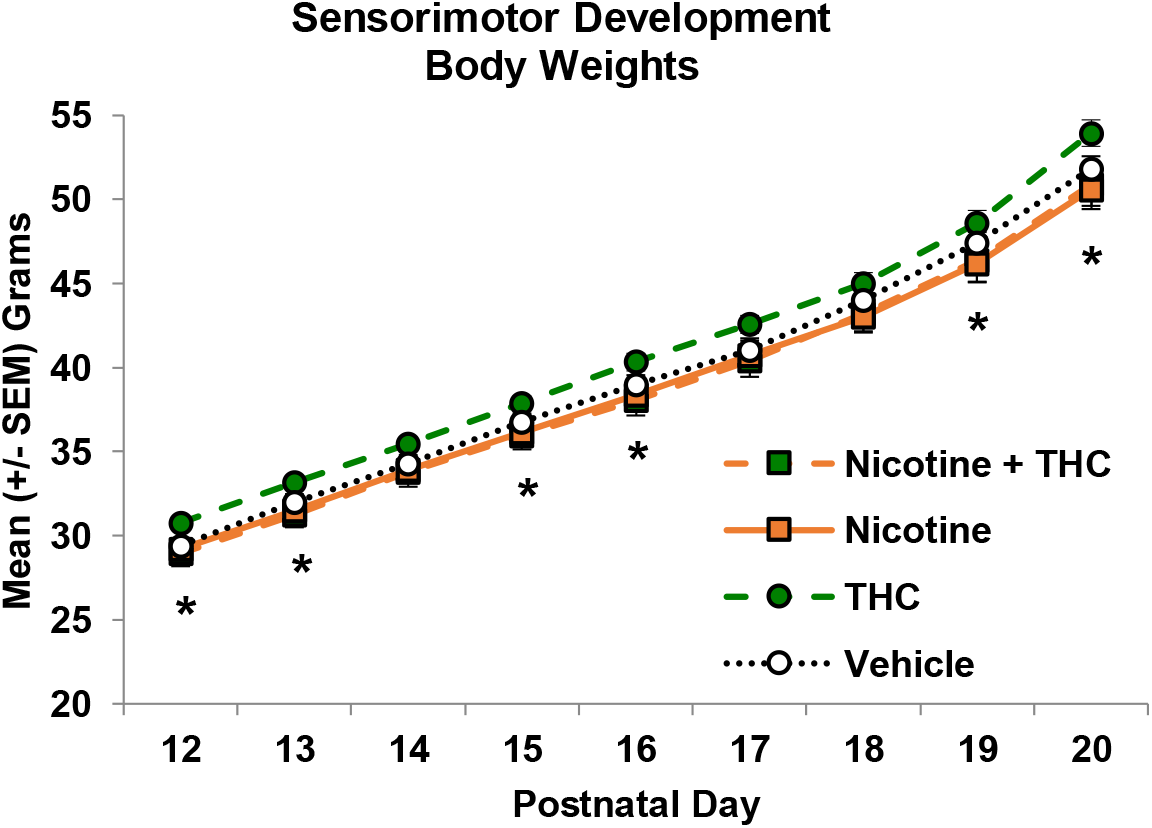
Increased weight among subjects exposed to THC alone drove a main effect of nicotine, when groups were collapsed. However, there were no significant differences in group weight when comparing all four groups

* = Nicotine < no Nicotine, *p*’s < .0.05

### Grip Strength and Hindlimb Coordination

Prenatal THC exposure impaired performance on the grip strength and hindlimb coordination task, delaying the first day of success (U = 914.00, *p* = 0.06). Although individual group analyses did not yield a statistically significant effect (H = 5.60, *p* = 0.13), the main effect of THC was driven by the combined prenatal exposure group (Nicotine+THC; Figure 3a). Day-by-day analyses (Figure 3b) indicated that fewer offspring exposed to combined Nicotine+THC prenatally were successful on PD 12 (Nicotine+THC < Nicotine: *X*^2^[1,48] = 3.00, *p* = 0.07; Nicotine+THC < Vehicle: *X*^2^[1,50] = 2.89, *p* = 0.07), effects that reached statistical significance by PD 13 (Nicotine+THC < THC: *X*^2^[1,46] = 7.33, *p* < 0.01; Nicotine+THC < Vehicle: *X*^2^[1,50] = 6.46, *p* < 0.05). Prenatal nicotine exposure alone did not alter any behavioral outcome variables in this task.

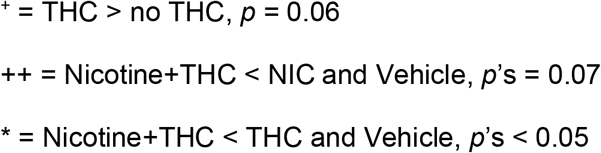

**Fig. 3.**
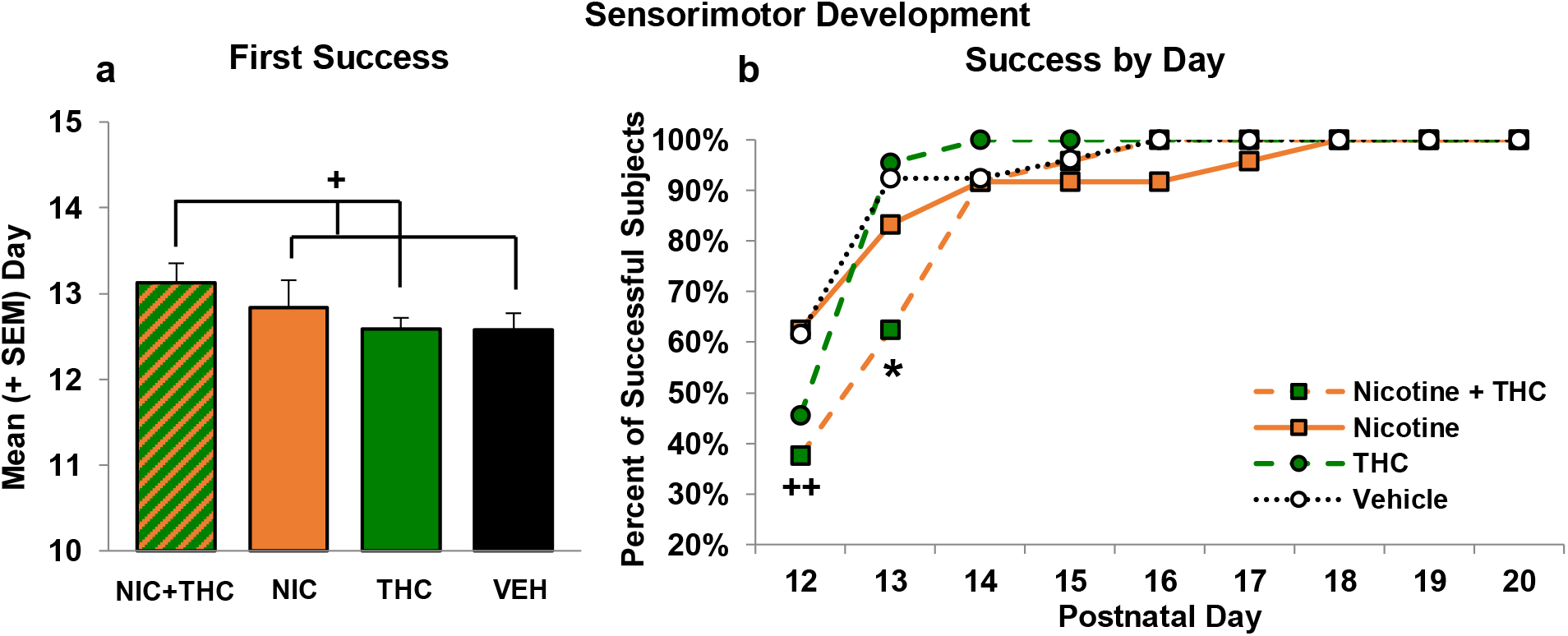
Combined exposure to nicotine and THC during gestation delayed successful performance on the early sensorimotor task, seen on the first day of success (a) and in the percent of subjects successful each day (b)

### Adolescent Parallel Bar Motor Coordination

#### Body Weights

By adolescence, females weighed less than males (F[1,65] = 52.04, *p* < 0.001) and gained weight more slowly compared to the males over the 3 days of parallel bar testing (Figure 4), leading to a Day*Sex interaction (F[2,130] = 51.91, *p* < 0.001) and main effect of Day (F[2,130] = 1418.78, *p* < 0.001).There were no significant main or interactive effects of prenatal nicotine or THC exposure on body weight.

**Fig. 4.**
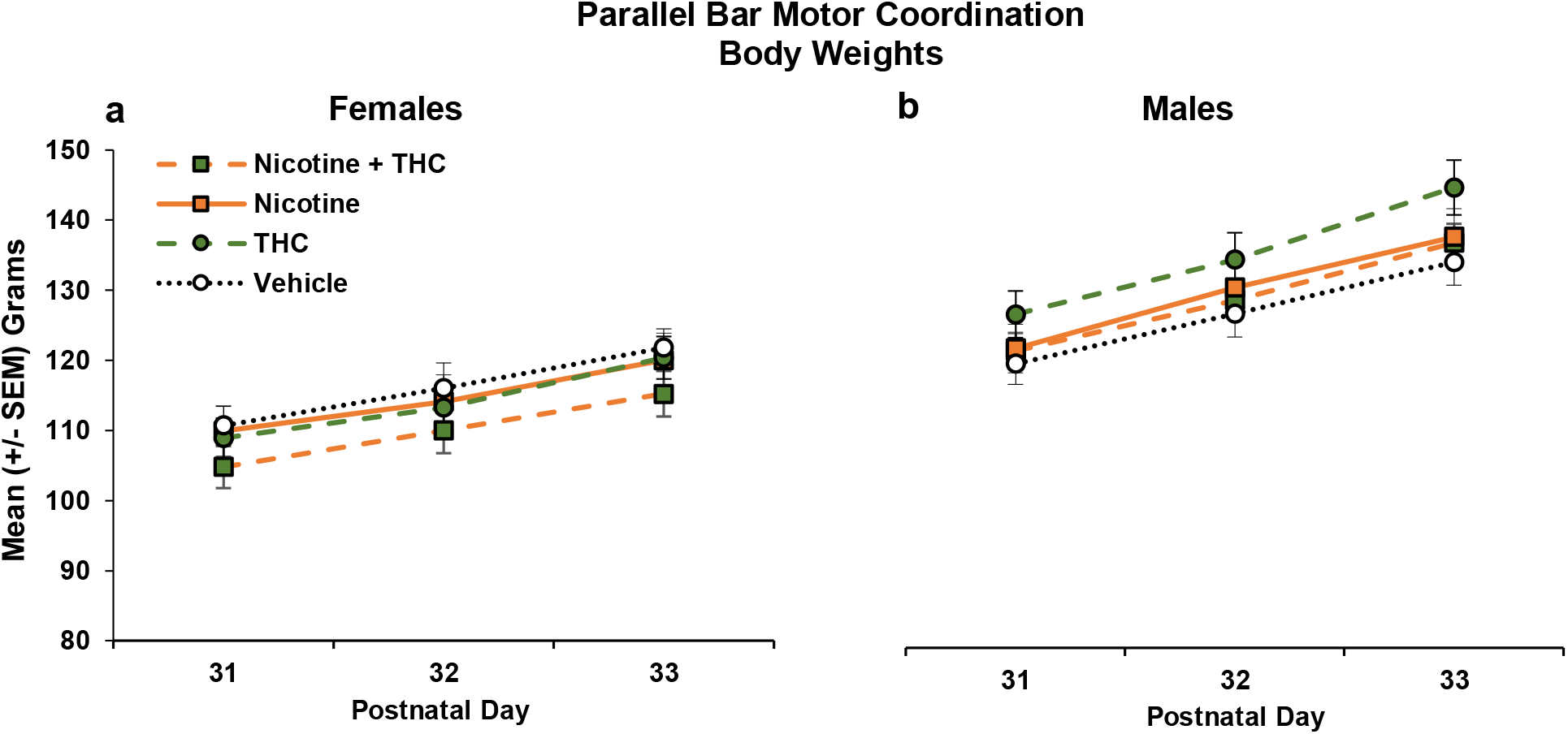
Females in all groups weighed significantly less (a) compared to males (b) during the parallel bar motor coordination task, but neither nicotine nor THC significantly affected body growth

#### Parallel Bar Motor Coordination

Some subjects failed to move during the motor coordination task and were, therefore, excluded from analyses. Importantly, these subjects represented all prenatal exposure groups (Nicotine+THC: 0 females, 1 male; Nicotine: 3 females, 0 males; THC: 3 females, 1 male; Vehicle: 2 females, 3 males).

##### Trials to First Success

There was a trend of a main effect of THC on the number of trials required for the first success on the parallel bar task (F[1,65] = 3.20, *p* = 0.08). Follow-up analyses comparing all groups (F[3,65] = 2.51, p = .07) indicated that this trend was driven by the combination exposure group (Nicotine+THC; Figure 5a). Post hoc analyses showed that subjects exposed to Nicotine+THC required significantly more trials to reach their first success compared to all other groups (LSD: *p*’s < 0.05). Although main or interactive effects of sex failed to reach statistical significance, this group effect was driven by females. Among females, there was a main effect of nicotine (F[1,32] = 4.18, *p* < 0.05) and an interaction of NIC*THC (F[1,32] = 3.97, *p* = 0.05). Post hoc analyses illustrated that the combination group required more trials to their first success compared to all other groups (F[3,32] = 3.51, *p* < 0.05; *p*’s <0.05; Figure 5b). In contrast, no main or interactive effects of prenatal nicotine or THC were observed among male offspring (Figure 5c).

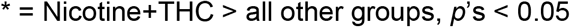

**Fig. 5.**
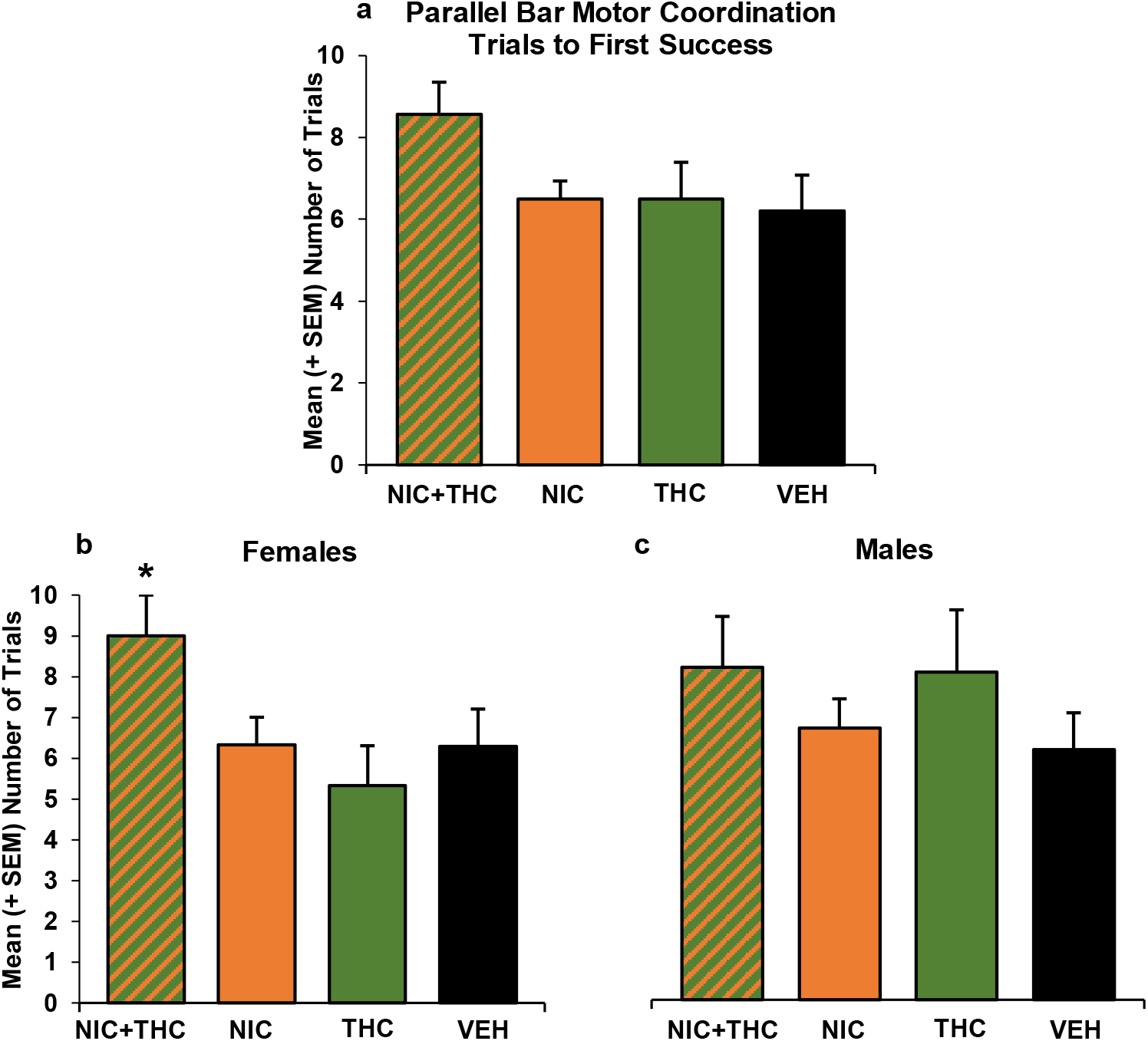
Subjects that were exposed to the combination of nicotine and THC required more trials to reach their first success compared to the other groups (a), an effect driven by female offspring (b, c)

##### Success Ratio

The success ratio for individual and total days was calculated by dividing the number of successful trials by total trials attempted. Across all testing days, subjects that were prenatally exposed to nicotine had fewer successes / total trials compared to subjects that were not exposed to nicotine (F[1,65] = 26.45, *p* < 0.001; Figure 6a). Moreover, each nicotine group performed significantly worse than the THC alone and the Vehicle control groups (all *p*’s < 0.05). On Day 1 of testing, there were no significant group differences in performance (Figure 6b). However, by Day 2, there were main effects of both Nicotine (F[1,69] = 12.57, *p* < 0.001) and THC (F[1,69] = 11.29, *p* < 0.002; Figure 6c). Group analyses confirmed that offspring exposed to either Nicotine or THC had lower success ratios compared to the Vehicle controls and that the combination impaired performance significantly more than all groups (F[3,65] = 8.07, *p* < 0.001). Notably, by day 3, there were group differences in starting widths; thus, task difficulty varied among groups, so one must be cautious of interpretations. Nevertheless, offspring exposed to Nicotine prenatally continued to have lower success ratios (F[1,65] = 6.91, *p* < 0.05; Figure 6d). Patterns were similar in males and females.

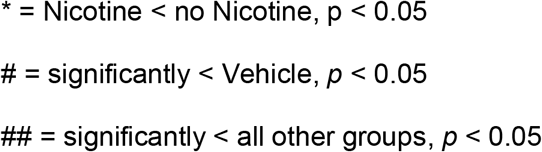

**Fig. 6.**
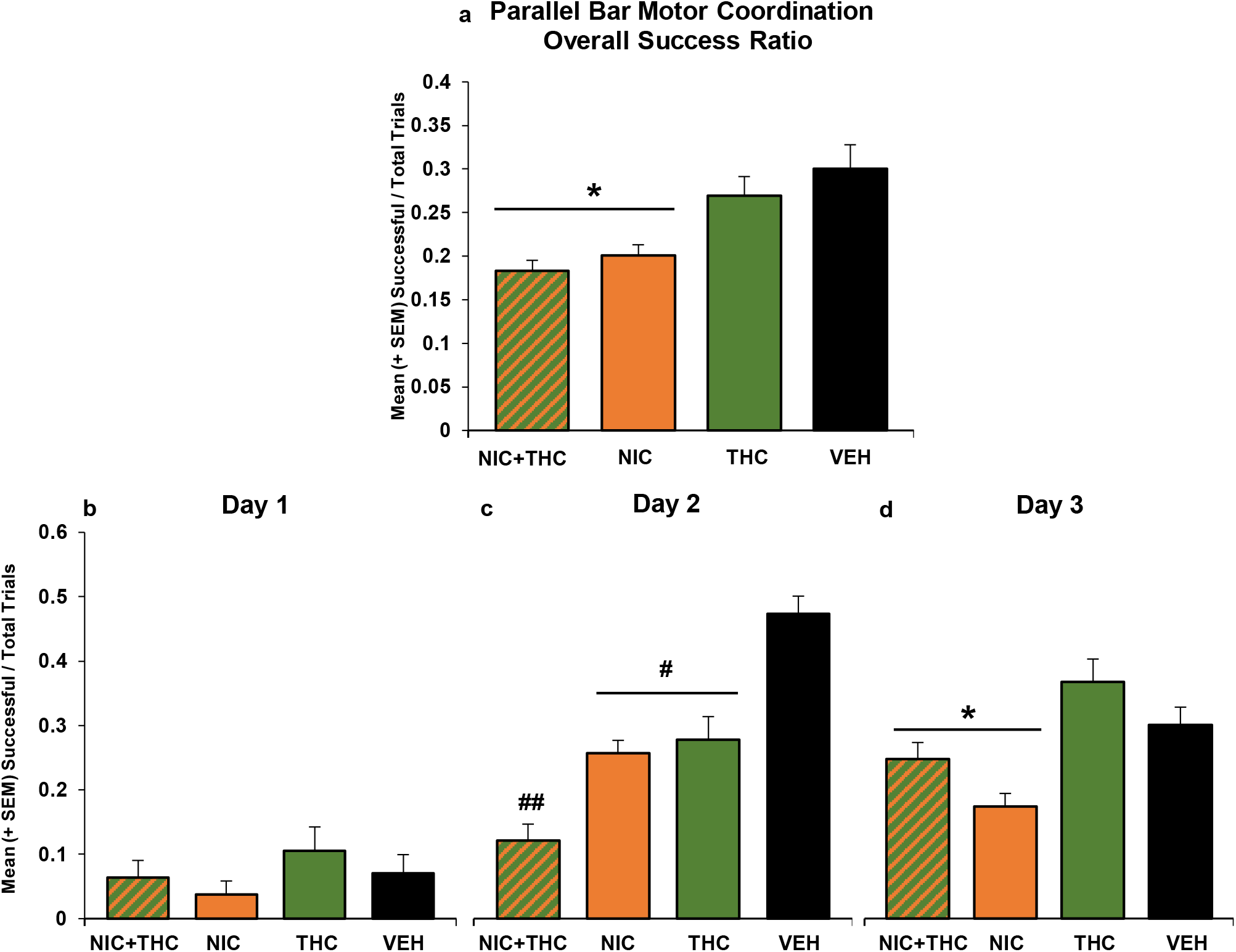
Subjects that were prenatally exposed to nicotine had lower success ratios across parallel bar testing (a). By Day 2 of testing, subjects that were prenatally exposed to nicotine or THC had lower success ratios compared to controls, effects that were additive in the combined group (c). By Day 3 (d), subjects exposed to prenatal nicotine continued to be less successful.

#### Maximum Width Achieved

A similar pattern was seen on the maximum rod width successfully traversed. Subjects’ performance improved over days (F[2,130] = 237.19, *p* < 0.001); however, both THC and nicotine impaired performance and the combination produced more severe effects, producing a main effect of THC (F[1,65] = 3.7, *p* < 0.05), a trending main effect of Nicotine (F[1,65] = 3.4, *p* = 0.06; Figure 7a), and an interaction of Day*THC (F[2,130] = 13.7, *p* < 0.001). Follow-up analyses confirmed a Day*Group interaction (F[6,130] = 5.43, *p* < 0.001).

**Fig. 7.**
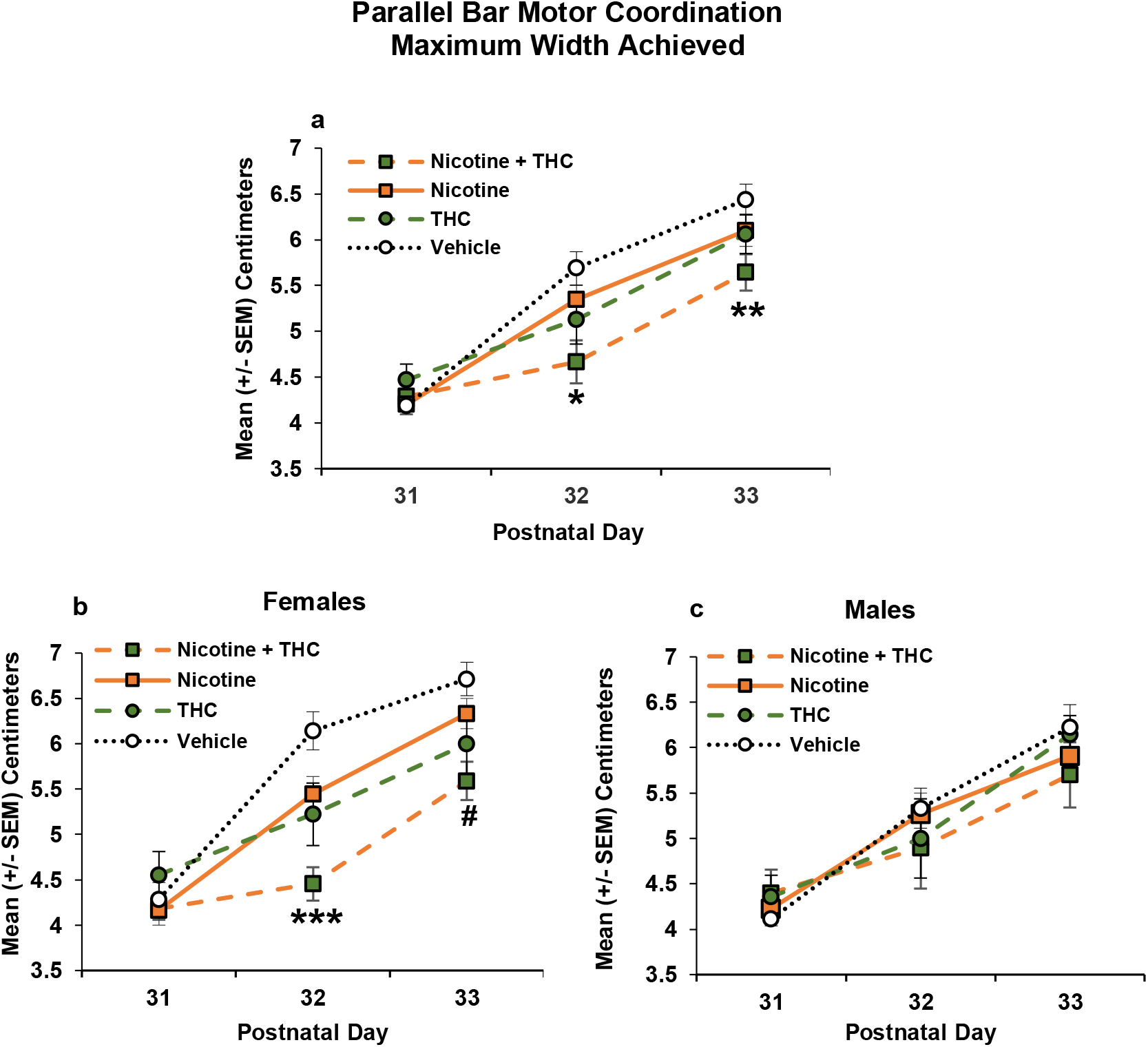
Offspring exposed prenatally to nicotine or THC achieved a lower maximum width during the motor coordination task, with the combination exposure group showing the greatest impairment (a). This effect was driven by the female offspring (b), whereas males showed no significant exposure effects (c)

There were no significant group differences in rod width successfully traversed on the first day of testing. However, by Day 2, subjects exposed to THC performed significantly worse compared to the subjects who were not prenatally exposed to THC (F[1,65] = 9.42, *p* < 0.01). Follow-up comparisons of all groups illustrated that the Combination group performed significantly worse than all other groups (Group: F[3,65] = 4.50, *p* < 0.05; p’s < 0.05). On Day 3, prenatal THC (F[1,65] = 5.02, *p* < 0.05) and Nicotine (F[1,65] = 3.94, *p* = 0.05) exposure impaired performance. Follow-up comparisons among all groups illustrated that only the combination exposure group differed significantly from vehicle controls (Group: F[3,65] = 3.12, *p* < 0.05). Although there were no significant interactions of sex any treatments, this effect was driven by the female offspring (Figure 7b, 7c).

Among female offspring, main effects of THC (F[1,32] = 8.00, *p <* 0.01), Nicotine (F[1,32] = 6.37, *p* < 0.05) and Day were found (F[2,64] = 120.42, *p* < 0.001), as well as interactions of Day*THC (F[2,64] = 11.63, *p* < 0.001). Analyses with all four groups confirmed a Day*Group interaction (F[6,64] = 4.5, *p* < 0.001). By Day 2, females exposed prenatally to THC (F[1,32] = 15.19, *p* < 0.001) or Nicotine (F[1,32] = 8.95, *p* < 0.01) successfully traversed a smaller maximum width. Importantly, these effects were additive, with either drug impairing performance compared to controls, and the combination of Nicotine+THC producing more severe deficits than either drug alone (Group: F[3,32] 8.14, *p* < 0.001; p’s < .05). By Day 3, females exposed to prenatal THC (F[1,32] = 8.60, *p* < 0.01) achieved a lower maximum width, an effect again driven by the combination group (Group: F[3,32] 3.74, *p* < 0.05, Nicotine+THC < Nicotine and Vehicle, *p*’s < 0.05).

In contrast, among males, although there was a significant interaction of Day*THC F[2,66] = 3.19, *p* = 0.05) and main effect of Day (F[2,66] = 116.77, *p* < 0.001), post hoc analyses did not indicate any significant differences among prenatal exposure groups on individual Days.

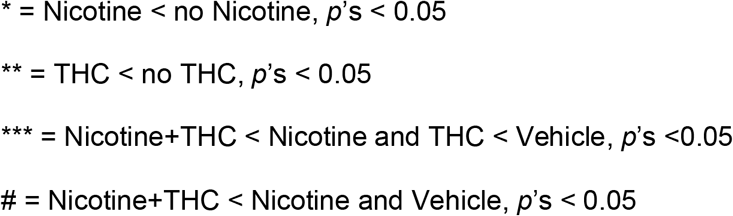

## DISCUSSION

The current study illustrates that prenatal exposure to either nicotine or THC impairs motor function, whereas the combination produces even more severe impairments. In this study, neither drug alone significantly affected early motor development, whereas combined exposure to prenatal nicotine and THC significantly delayed sensorimotor development. In contrast, prenatal exposure to either nicotine or THC impaired performance on the parallel bar motor coordination task during early adolescence, and combined exposure exacerbated these effects.

Although there is a rich literature on the effects of nicotine on development, clinical and preclinical research examining the long-term effects of prenatal nicotine exposure via e-cigarettes is limited, especially those investigating its impacts on behavioral development. Numerous studies have reported that prenatal nicotine exposure via traditional cigarettes impairs motor function, including deficits in motor control and coordination (Cornelius et al., 2001; Ernst et al., 2001; Larsson & Montgomery, 2011; Römer et al., 2020). Importantly, a more recent study found that nicotine consumed via e-cigarettes has similar consequences to traditional cigarettes on infant neurobehavior, leading to abnormal reflexes and deficits in motor maturity, including crawling, standing, and steps (Froggatt et al., 2020). In that study, motor dysfunction was also positively correlated with the number of years a mother smoked prior to conception, as well as the dose consumed (Froggatt et al., 2020).

Preclinical studies have also found that prenatal nicotine exposure using combustible cigarettes, nicotine pumps, i.p or s.c. injections, can alter offspring motor development, delaying righting reflexes, increasing motor activity, and impairing motor learning skills (Ajarem & Ahmad, 1998; Blood-Siegfried & Rende, 2010; Lee & Picciotto, 2019; Willford et al., 2010). However, this is the first study, to date, to examine motor development following nicotine exposure via e-cigarettes in pregnant dams. Although we did not find evidence that prenatal nicotine exposure via e-cigarettes impaired early sensorimotor development, we did find that prenatal e-cigarette exposure led to motor deficits during adolescence. The nicotine dose used in this study (36 mg/mL) produced maternal plasma levels of ~20 ng/mL (Breit et al., 2021). This dose reflects nicotine concentrations used by moderate-heavy smokers (Breland et al., 2017), and the plasma levels also model those observed in human populations, as well as animal models system, of moderate-high doses via various administrative routes, including e-cigarettes (Farsalinos & Polosa, 2014; Farsalinos et al., 2015; Matta et al., 2007; Montanari et al., 2020). Thus, although e-cigarettes are assumed to be a safer route for nicotine consumption (Mark et al., 2015; McCubbin et al., 2017), these results suggest that prenatal nicotine exposure via e-cigarettes may lead to motor deficits, as observed with other administrative routes.

Consistent with motor impairments, prenatal nicotine exposure may lead to neuropathology in CNS areas important for motor function (Bruin et al., 2010; Dwyer et al., 2009). Prenatal nicotine exposure can directly affect nicotinic acetylcholine receptors (nAChRs), as nicotine crosses the placenta and binds to fetal nAChRs, which are present during early neural tube development (Blood-Siegfried & Rende, 2010; Wickstrom, 2007). Importantly, motor areas of the brain such as the basal ganglia and cerebellum are rich in nAChRs (Blood-Siegfried & Rende, 2010; Graham et al., 2002; Wickstrom, 2007). Thus, these CNS areas may be particularly vulnerable to prenatal nicotine exposure. For example, animal studies have shown that prenatal nicotine exposure can change the subunit composition of nAChRs (Lv et al., 2008; Turner, 2005) and increase sensitivity of nAChRs receptors in areas such as the cortex and cerebellum (Lv et al., 2008; Slotkin et al., 1999). Similarly, fetal human brain tissue exposed to cigarette smoke during development shows increased alpha 4 nAChR subunit expression in the brainstem and cerebellum as early as the first trimester (Falk el at., 2005). Thus, changes to nAChRs during fetal development, which are important for neural circuity development in both the central and peripheral nervous systems (Dwyer et al., 2008), may adversely affect brain and behavioral development.

Similarly, in the current study, prenatal exposure to THC via an e-cigarette did not significantly influence sensorimotor development on its own (although a trend was present), but did lead to long-lasting deficits in motor coordination during early adolescence. We previously reported that prenatal exposure to THC (with or without combined alcohol) delays sensorimotor development, but like the present study, impairs motor coordination in adolescence on the parallel bar task (Breit et al., 2021). Previous studies examining the consequences of prenatal cannabis on motor development have been mixed. Prenatal cannabis extract exposure has been shown to impair rotarod performance (Abel, 1979) and lead to hyperactivity (Campolongo et al., 2011); however, other studies have failed to find or fails to impact motor behaviors (Abel, 1976; Vardaris et al., 1976). Importantly, the impact of developmental cannabinoid exposure may largely depend on the timing of exposure, type of cannabinoid used, dose used, and/or the route of administration. For example, we have also previously shown that exposure to high levels of synthetic cannabinoids (CP-55,940; i.p.) during the early postnatal brain growth spurt (PD 4-9) actually advance the trajectory of early sensorimotor development, but have no effect on adolescent motor coordination (Breit et al., 2019).

Clinical studies examining the effects of prenatal cannabis exposure on motor development are limited and have similarly yielded mixed results (Grant et al., 2018; Metz & Borgelt, 2018). For example, prenatal cannabis exposure may lead to increased startles and tremors in neonates and advancement of motor development in toddlers (Fried et al., 1998); however, some studies have failed to find any alterations in balance or motor coordination (Chandler et al., 1996). Importantly, these studies are limited, and the prenatal exposure of these subjects occurred decades ago; thus, results will likely not generalize to the cannabis products and consumption patterns of today. In the current study, the peak levels of THC mimic those from low-moderate THC level products used today (Andrenyak et al., 2017; Nguyen et al., 2016), with maternal plasma THC levels (15-30 mins post-exposure) of 20 ng/mL. It is alarming that such levels produce behavioral effects on the offspring, given that potency levels of THC in cannabis products are consistently rising (ElSohly et al., 2021).

The endocannabinoid system plays an important role during pregnancy (Ezechukwu et al., 2020; Metz & Borgelt, 2018). For example, endocannabinoids are important for placenta development, which expresses both CB1 and CB2 receptors (Habayeb et al., 2008); thus, cannabis consumption could influence placental function. Importantly, the cerebellum, an area critical for motor function, also has a high density of CB1 receptors (Herkenham et al., 1991; Roncero et al., 2020; Takahashi & Linden, 2000). Cerebellar CB1 receptors play a role in early synaptic communication in rodents as early as GD 11-14 (Berghuis et al., 2007; Harkany et al., 2007), overlapping with the exposure period of the current study (GD 5-20). However, the timing of exposure is important to consider. Importantly, if cannabinoid exposure occurs during the early postnatal period in rodents (PD 0-14), it may selectively target CB1 binding in GABAergic neurons in the cerebellum (Diana et al., 2002), basal ganglia, and cortex (Berrendero et al., 1999), which would up-regulate GABA activity and enhance motor performance (Benagiano et al., 2007; Chiu et al., 2005). Developmental changes in the endocannabinoid system within the cerebellum may account for the inconsistent motor effects reported in the prenatal cannabis literature, and again highlights the importance of considering exposure timing, type, dose, and route in this field.

Importantly, the combination of nicotine and THC via e-cigarettes produced more severe motor impairments than either drug alone. Combined prenatal exposure to nicotine and THC via e-cigarettes delayed early sensorimotor development, a synergistic effect as neither nicotine nor THC by itself significantly affected performance on this task. In contrast, the effects of combined exposure on long-lasting deficits in motor coordination were additive. To our knowledge, this is the first study to examine behavioral effects of prenatal co-exposure of nicotine and THC exposure via an e-cigarette. These results are intriguing, given that the combination of nicotine and THC reduces plasma levels of both nicotine and THC, particularly early in gestation (Breit et al., 2021). Clinical studies of prenatal nicotine/tobacco and cannabis exposure are similarly limited. However, similar to our findings, prenatal co-exposure to tobacco and cannabis produces more severe alterations in infants than exposure to either drug alone, producing reductions in self-soothing, attention deficits, decreased motor activity (Stroud et al., 2018), reduced cortisol responses (Eiden et al., 2020), and poorer autonomic regulation (Eiden et al., 2018). Motor-related domains have not yet been investigated in clinical populations. Importantly, given the ease of co-use in e-cigarettes and given that the drug levels used in the current study are ecologically valid, it is concerning that the combination may be more damaging to the fetus.

Interestingly, although there were no effects of sex on early sensorimotor development, the consequences of prenatal nicotine and/or THC exposure on parallel bar performance were more robust among female offspring. For example, even though success ratios were similar between sexes, the combined exposure significantly increased the number of trials required for a success among females, but not males. Similarly, maximum width successfully traversed was impaired by either nicotine or THC, with additive effects of combined exposure among females, but not males. It is important to note that the width between the parallel bars expands by 0.5-cm with each successful trial; thus, sex-related differences in body size may have played a role on performance for this measure. However, we found no significant relationship between body weight and successful performance. We did not find sex differences in this task in our previous work examining prenatal THC exposure (Breit et al., 2021), however, other have reported more severe motor impairments in female offspring (Abel, 1979).

Reports of sex-dependent effects of prenatal exposure to these drugs have been mixed. For example, males exposed to prenatal nicotine exposure have exhibited greater alterations in a pellet-reaching task (Lee & Picciotto, 2019), rearing activity (Peters & Tang, 1982), locomotor/stereotypy activity (Shacka et al., 1997), and anxiety-related behaviors (Polli et al., 2020). In contrast, although it has been reported that females are more affected by prenatal nicotine exposure on learning tasks (Levin et al., 1996), others fail to find sex differences following prenatal nicotine exposure in domains such as activity level (Polli et al., 2020) and maze learning (Sorenson et al., 1991). Although data are limited, several studies suggest that males may be more vulnerable to prenatal THC exposure than females in domains such as social interaction (Bara et al., 2018), ultrasonic vocalizations, and hyperactivity (Manduca et al., 2020), but that sex effects vary widely on learning and anxiety measures (Keeley et al., 2015). Regardless, given the limited number of studies examining prenatal co-exposure to these drugs, it is difficult to understand the mechanism of the sex differences observed in the current study, which future studies should aim to unfold.

There are several limitations to note in this study. First, the control group in this study was exposed to the vehicle (propylene glycol) prenatally, and it is still unknown whether propylene glycol, itself, may exert teratogenic effects. However, we did complete a small pilot study that compared the vehicle group to a group of non-handled, non-exposed controls and we found no differences in motor performance between these groups. Importantly, prenatal exposure to nicotine and/or THC led impaired motor development significantly more than prenatal vehicle exposure. Secondly, this study only included one dose of each drug. Even with these single doses we found both additive and synergistic effects, but a more extensive examination of multiple doses would be valuable. Lastly, this study focused on nicotine and THC only. Although e-cigarettes allow for consumption of these isolated compounds, products on the market vary in terms of the constituents.

In conclusion, we found that prenatal exposure to nicotine and/or THC via an e-cigarette model leads to deficits in motor development, and the combination of nicotine and THC exposure produces more severe impairments. These results suggest that co-consumption of nicotine and cannabis via e-cigarettes may not be as safe as assumed, and may lead to long-lasting alterations in motor-related domains among offspring. These data have important implications for pregnant women, public health, and public policy decisions regarding prenatal e-cigarette use.

## ACKNOWLEDGMENTS

This work was supported by Tobacco-Related Disease Research Program grant (28IP-0026) to JDT. It was also supported by a National Institute on Alcohol Abuse and Alcoholism training grant (T32AA007456-38) and a National Institutes of Health Loan Repayment Program award to KRB. THC was obtained from the National Institute on Drug Abuse Drug Supply Program. Special thanks to Maury Cole at La Jolla Alcohol Research, Inc. (San Diego, CA) for building the vapor inhalation equipment and for co-exposure advisement. Lastly, we want to recognize the members of the Center for Behavioral Teratology at San Diego State University for assisting in data collection and interpretation, particularly the instrumental efforts of Cristina Rodriguez, Karen Thomas, Jaclyn Hanson, and Jacob Ruiz.

## REFERENCES

Abel, E. (1979). Behavioral teratology of marihuana extract in rats. Neurobehav. Toxicol, 1(4), 285–287.

Abel, E. L. (1976). Clinical and epidemiological studies NIDA Res Monogr, 59, 20–35.

Ajarem, J. S., & Ahmad, M. (1998). Prenatal nicotine exposure modifies behavior of mice through early development Pharmacol Biochem Behav., 59(2), 313–318. https://doi.org/10.1016/s0091-3057(97)00408-5

Amlani, A., Hornick, M. G., Cooper, K., Prazad, P., Donovan, R., & Gulati, A. (2017). Maternal cannabinoid use alters cannabinoid (cb1) and endothelin (etb) receptor expression in the brains of dams but not their offspring. Dev Neurosci, 39(6), 498–506. https://doi.org/10.1159/000480453

Andrenyak, D. M., Moody, D. E., Slawson, M. H., O’Leary, D. S., & Haney, M. (2017). Determination of∆-9-tetrahydrocannabinol (thc), 11-hydroxy-thc, 11-nor-9-carboxy-thc and cannabidiol in human plasma using gas chromatography–tandem mass spectrometry. J. Anal. Toxicol., 41(4), 277–288. https://doi.org/10.1093/jat/bkw136

Bara, A., Manduca, A., Bernabeu, A., Borsoi, M., Serviado, M., Lassalle, O., Murphy, M., Wager-Miller, J., Mackie, K., Pelissier-Alicot, A.-L., Trezza, V., & Manzoni, O. J. (2018). Sex-dependent effects of in utero cannabinoid exposure on cortical function. ELife, 7, 1–31. https://doi.org/10.7554/elife.36234

Benagiano, V., Lorusso, L., Flace, P., Girolamo, F., Rizzi, A., Sabatini, R., Auteri, P., Bosco, L., Cagiano, R., & Ambrosi, G. (2007). Effects of prenatal exposure to the cb-1 receptor agonist win 55212-2 or co on the gabaergic neuronal systems of rat cerebellar cortex. Neurosci., 149(3), 592–601. https://doi.org/10.1016/j.neuroscience.2007.07.050

Berghuis, P., Rajnicek, A. M., Morozov, Y. M., Ross, R. A., Mulder, J., Urbán, G. M., Monory, K., Marsicano, G., Matteoli, M., & Canty, A. (2007). Hardwiring the brain: Endocannabinoids shape neuronal connectivity. Science, 316(5828), 1212–1216. https://doi.org/10.1126/science.1137406

Berrendero, F., Sepe, N., Ramos, J. A., Di Marzo, V., & Fernández-Ruiz, J. J. (1999). Analysis of cannabinoid receptor binding and mrna expression and endogenous cannabinoid contents in the developing rat brain during late gestation and early postnatal period. Synapse, 33(3), 181–191. https://doi.org/10.1002/(SICI)1098-2396(19990901)33:3

Blood-Siegfried, J., & Rende, E. K. (2010). The long-term effects of prenatal nicotine exposure on neurologic development. Journal of Midwifery & Women’s Health, 55(2), 143–152. https://doi.org/10.1016/j.jmwh.2009.05.006

Breit, K. R., Rodriguez, C., Hussain, S., Thomas, K., Zeigler, M., Gerasimidis, I., & Thomas, J. (2021). Pharmacokinetic effects of combined exposure to nicotine and thc via e-cigarettes in pregnant rats. BioRxiv. https://doi.org/10.1101/2021.07.07.451537

Breit, K. R., Rodriguez, C., Lei, A., Hussain, S., & Thomas, J. D. (2021). Prenatal alcohol and thc e-cigarette exposure effects on motor development. BioRxiv. https://doi.org/10.1101/2021.05.26.445823

Breit, K. R., Zamudio, B., & Thomas, J. D. (2019). Altered motor development following late gestational alcohol and cannabinoid exposure in rats. Neurotoxicol Teratol, 73, 31–41. https://doi.org/10.1016/j.ntt.2019.03.005

Breland, A., Soule, E., Lopez, A., Ramôa, C., El-Hellani, A., & Eissenberg, T. (2017). Electronic cigarettes: What are they and what do they do? Ann. N. Y. Acad. Sci., 1394(1), 1–40.

Brown, Q. L., Sarvet, A. L., Shmulewitz, D., Martins, S. S., Wall, M. M., & Hasin, D. S. (2017). Trends in marijuana use among pregnant and nonpregnant reproductive-aged women, 2002–2014. JAMA, 317(2), 207–2011. https://doi.org/10.1001/jama.2016.17383

Bruin, J. E., Gerstein, H. C., & Holloway, A. C. (2010). Long-term consequences of fetal and neonatal nicotine exposure: A critical review. Toxicol. Sci., 116(2), 364–374. https://doi.org/10.1093/toxsci/kfq103

Campolongo, P., Trezza, V., Ratano, P., Palmery, M., & Cuomo, V. (2011). Developmental consequences of perinatal cannabis exposure: Behavioral and neuroendocrine effects in adult rodents. Psychopharmacology, 214(1), 5–15. https://doi.org/10.1007/s00213-010-1892-x

Chandler, L. S., Richardson, G. A., Gallagher, J. D., & Day, N. L. (1996). Prenatal exposure to alcohol and marijuana: Effects on motor development of preschool children. Alcohol Clin Exp Res, 20(3), 455–461. https://doi.org/10.1111/j.1530-0277.1996.tb01075.x

Chandra, S., Radwan, M. M., Majumdar, C. G., Church, J. C., Freeman, T. P., & Elsohly, M. A. (2019). New trends in cannabis potency in USA and europe during the last decade (2008–2017). Eur Arch Psychiatry Clin Neurosci . 269(1), 5–15. https://doi.org/10.1007/s00406-019-00983-5

Chiu, C.-S., Brickley, S., Jensen, K., Southwell, A., Mckinney, S., Cull-Candy, S., Mody, I., & Lester, H. A. (2005). Gaba transporter deficiency causes tremor, ataxia, nervousness, and increased gaba-induced tonic conductance in cerebellum. J. Neurosci, 25(12), 3234–3245. https://doi.org/10.1523/JNEUROSCI.3364-04.2005

Coleman-Cowger, V. H., Oga, E. A., Peters, E. N., & Mark, K. (2018). Prevalence and associated birth outcomes of co-use of cannabis and tobacco cigarettes during pregnancy. Neurotoxicol Teratol, 68, 84–90. https://doi.org/10.1016/j.ntt.2018.06.001

Coleman-Cowger, V. H., Schauer, G. L., & Peters, E. N. (2017). Marijuana and tobacco co-use among a nationally representative sample of us pregnant and non-pregnant women: 2005–2014 national survey on drug use and health findings. Drug Alcohol Depend., 177, 130–135. https://doi.org/10.1016/j.drugalcdep.2017.03.025

Cornelius, M. D., Ryan, C. M., Day, N. L., Goldschmidt, L., & Willford, J. A. (2001). Prenatal tobacco effects on neuropsychological outcomes among preadolescents. J. Dev. Behav. Pediatr., 22(4), 217–225. https://doi.org/10.1097/00004703-200108000-00002

Dahl, R. E., Scher, M. S., Williamson, D. E., Robles, N., & Day, N. (1995). A longitudinal study of prenatal marijuana use. Effects on sleep and arousal at age 3 years. Arch Pediatr Adolesc Med, 149(2), 145–150. https://doi.org/10.1001/archpedi.1995.02170140027004

Diana, M. A., Levenes, C., Mackie, K., & Marty, A. (2002). Short-term retrograde inhibition of gabaergic synaptic currents in rat purkinje cells is mediated by endogenous cannabinoids. J. Neurosci, 22(1), 200–208. https://doi.org/10.1523/jneurosci.22-01-00200.2002

Dwyer, J. B., Broide, R. S., & Leslie, F. M. (2008). Nicotine and brain development. Defects Res. C, 84(1), 30–44. https://doi.org/10.1002/bdrc.20118

Dwyer, J. B., McQuown, S. C., & Leslie, F. M. (2009). The dynamic effects of nicotine on the developing brain. Pharmacol. Ther., 122(2), 125–139. https://doi.org/10.1016/j.pharmthera.2009.02.003

Eiden, R. D., Schuetze, P., Shisler, S., & Huestis, M. A. (2018). Prenatal exposure to tobacco and cannabis: Effects on autonomic and emotion regulation. Neurotoxicol Teratol, 68, 47–56. https://doi.org/10.1016/j.ntt.2018.04.007

Eiden, R. D., Shisler, S., Granger, D. A., Schuetze, P., Colangelo, J., & Huestis, M. A. (2020). Prenatal tobacco and cannabis exposure: Associations with cortisol reactivity in early school age children. Int. J. Behav. Med, 27(3), 343–356. https://doi.org/10.1007/s12529-020-09875-8

ElSohly, M. A., Chandra, S., Radwan, M., Majumdar, C. G., & Church, J. C. (2021). A comprehensive review of cannabis potency in the united states in the last decade. Biol Psychiatry Cogn Neurosci Neuroimaging, 6(6), 603–606. https://doi.org/10.1016/j.bpsc.2020.12.016

Ernst, M., Moolchan, E. T., & Robinson, M. L. (2001). Behavioral and neural consequences of prenatal exposure to nicotine. J Am Acad Child Adolesc Psychiatry, 40(6), 630–641. https://doi.org/10.1097/00004583-200106000-00007

Ezechukwu, H. C., Diya, C. A., Shrestha, N., & Hryciw, D. H. (2020). Role for endocannabinoids in early pregnancy: Recent advances and the effects of cannabis use. Am. J. Physiol. Endocrinol. Metab. https://doi.org/10.1152/ajpendo.00210.2020

Farsalinos, K. E., & Polosa, R. (2014). Safety evaluation and risk assessment of electronic cigarettes as tobacco cigarette substitutes: A systematic review. Ther. Adv. Drug Saf., 5(2), 67–86. https://doi.org/10.1177/2042098614524430

Farsalinos, K. E., Spyrou, A., Stefopoulos, C., Tsimopoulou, K., Kourkoveli, P., Tsiapras, D., Kyrzopoulos, S., Poulas, K., & Voudris, V. (2015). Nicotine absorption from electronic cigarette use: Comparison between experienced consumers (vapers) and naïve users (smokers). Sci. Rep., 5(1), 1–9. https://doi.org/10.1038/srep11269

Fish, E., Boschen, K., Murdaugh, L., Mendoza-Romero, H., Williams, K., & Parnell, S. (2017). Alcohol exacerbates the teratogenic effects of prenatal cannabinoid exposure in a c57bl/6j mouse model. Birth Defects Research,

Fish, E., Gilbert, M., Sulik, K., & Parnell, S. (2016). Ethanol and the synthetic cannabinoid, cp-55,940, are synergistically teratogenic in a mouse model. Alcoholism-Clinical and Experimental Research,

Forray, A. (2016). Substance use during pregnancy. F1000Research, 5, 1–9. https://doi.org/10.12688/f1000research.7645.1

Fried, P. A. (1991). Marijuana use during pregnancy: Consequences for the offspring. Seminars in Perinatology 15(4), 280–287.

Fried, P. A. (1996). Behavioral outcomes in preschool and school-age children exposed prenatally to marijuana: A review and speculative interpretation. NIDA Res. Monogr., 164, 242–260.

Fried, P. A., Watkinson, B., & Gray, R. (1998). Differential effects on cognitive functioning in 9-to 12-year olds prenatally exposed to cigarettes and marihuana. Neurotoxicol Teratol, 20(3), 293–306.

Froggatt, S., Reissland, N., & Covey, J. (2020). The effects of prenatal cigarette and e-cigarette exposure on infant neurobehaviour: A comparison to a control group. EClinicalMedicine, 28, 1–6. https://doi.org/10.1016/j.eclinm.2020.100602

Glasser, A. M., Collins, L., Pearson, J. L., Abudayyeh, H., Niaura, R. S., Abrams, D. B., & Villanti, A. C. (2017). Overview of electronic nicotine delivery systems: A systematic review. Am. J. Prev. Med., 52(2), 1–63. https://doi.org/10.1016/j.amepre.2016.10.036

Goldschmidt, L., Richardson, G. A., Larkby, C., & Day, N. L. (2016). Early marijuana initiation: The link between prenatal marijuana exposure, early childhood behavior, and negative adult roles. Neurotoxicol Teratol, 58, 40–45. https://doi.org/10.1016/j.ntt.2016.05.011

Graham, A., Court, J. A., Martin-Ruiz, C. M., Jaros, E., Perry, R., Volsen, S. G., Bose, S., Evans, N., Ince, P., Kuryatov, A., Lindstrom, J., Gotti, C., & Perry, E. K. (2002). Immunohistochemical localisation of nicotinic acetylcholine receptor subunits in human cerebellum. Neurosci., 113(3), 493–507. https://doi.org/10.1016/s0306-4522(02)00223-3

Grant, K. S., Petroff, R., Isoherranen, N., Stella, N., & Burbacher, T. M. (2018). Cannabis use during pregnancy: Pharmacokinetics and effects on child development. Pharmacol. Ther., 182, 133–151. https://doi.org/10.1016/j.pharmthera.2017.08.014

Gunn, J. K. L., Rosales, C. B., Center, K. E., Nunez, A. V., Gibson, S. J., & Ehiri, J. E. (2015). The effects of prenatal cannabis exposure on fetal development and pregnancy outcomes: A protocol. BMJ Open, 5(3), 1–5. https://doi.org/10.1136/bmjopen-2014-007227

Habayeb, O. M. H., Taylor, A. H., Bell, S. C., Taylor, D. J., & Konje, J. C. (2008). Expression of the endocannabinoid system in human first trimester placenta and its role in trophoblast proliferation. Endocrinology, 149(10), 5052–5060. https://doi.org/10.1210/en.2007-1799

Hall, W., & Weier, M. (2015). Assessing the public health impacts of legalizing recreational cannabis use in the USA. Clin. Pharmacol. Ther., 97(6), 607–615. https://doi.org/10.1002/cpt.110

Harkany, T., Guzman, M., Galve-Roperh, I., Berghuis, P., Devi, L. A., & Mackie, K. (2007). The emerging functions of endocannabinoid signaling during cns development. Trends Pharmacol. Sci., 28(2), 83–92. https://doi.org/10.1016/j.tips.2006.12.004

Herkenham, M., Lynn, A., Johnson, M., Melvin, L., De Costa, B., & Rice, K. (1991). Characterization and localization of cannabinoid receptors in rat brain: A quantitative in vitro autoradiographic study. J. Neurosci, 11(2), 563–583. https://doi.org/10.1523/jneurosci.11-02-00563.1991

Holbrook, B. D. (2016). The effects of nicotine on human fetal development. Defects Res. C, 108(2), 181–192. https://doi.org/10.1002/bdrc.21128

Holz, N. E., Boecker, R., Baumeister, S., Hohm, E., Zohsel, K., Buchmann, A. F., Blomeyer, D., Jennen-Steinmetz, C., Hohmann, S., Wolf, I., Plichta, M. M., Meyer-Lindenberg, A., Banaschewski, T., Brandeis, D., & Laucht, M. (2014). Effect of prenatal exposure to tobacco smoke on inhibitory control. Am. J. Psychiatry, 71(7), 786–796. https://doi.org/10.1001/jamapsychiatry.2014.343

Huizink, A. C. (2014). Prenatal cannabis exposure and infant outcomes: Overview of studies. Prog Neuropsychopharmacol Biol Psychiatry, 52, 45–52. https://doi.org/10.1016/j.pnpbp.2013.09.014

Hunter, A., Murray, R., Asher, L., & Leonardi-Bee, J. (2020). The effects of tobacco smoking, and prenatal tobacco smoke exposure, on risk of schizophrenia: A systematic review and meta-analysis. N&TR, 22(1), 3–10. https://doi.org/10.1093/ntr/nty160

Keeley, R., Trow, J., Bye, C., & McDonald, R. (2015). Part ii: Strain-and sex-specific effects of adolescent exposure to thc on adult brain and behaviour: Variants of learning, anxiety and volumetric estimates. Behav. Brain Res, 288, 132–152. https://doi.org/10.1016/j.bbr.2015.01.001

Lambers, D. S., & Clark, K. E. (1996). The maternal and fetal physiologic effects of nicotine. Semin. Perinatol., 20(2), 115–126. https://doi.org/10.1016/s0146-0005(96)80079-6

Larsson, M., & Montgomery, S. M. (2011). Maternal smoking during pregnancy and physical control and coordination among offspring. J Epidemiol Community Health, 65(12), 1151–1158. https://doi.org/10.1136/jech.2008.085241

Lee, A. M., & Picciotto, M. R. (2019). Perinatal nicotine exposure impairs learning of a skilled forelimb reaching task in male but not female adult mice. Behav. Brain Res, 367, 176–180. https://doi.org/10.1016/j.bbr.2019.04.007

Levin, E. D., Wilkerson, A., Jones, J. P., Christopher, N. C., & Briggs, S. J. (1996). Prenatal nicotine effects on memory in rats: Pharmacological and behavioral challenges. Brain Res. Dev. Brain Res., 97(2), 207–215. https://doi.org/10.1016/s0165-3806(96)00144-7

Longo, C., Fried, P., Cameron, I., & Smith, A. (2014). The long-term effects of prenatal nicotine exposure on verbal working memory: An fmri study of young adults. Drug Alcohol Depend, 144, 61–69. https://doi.org/10.1016/j.drugalcdep.2014.08.006

Lv, J., Mao, C., Zhu, L., Zhang, H., Pengpeng, H., Xu, F., Liu, Y., Zhang, L., & Xu, Z. (2008). The effect of prenatal nicotine on expression of nicotine receptor subunits in the fetal brain. NeuroToxicology, 29(4), 722–726. https://doi.org/10.1016/j.neuro.2008.04.015

Manduca, A., Servadio, M., Melancia, F., Schiavi, S., Manzoni, O. J., & Trezza, V. (2020). Sex-specific behavioural deficits induced at early life by prenatal exposure to the cannabinoid receptor agonist win55, 212-2 depend on mglu5 receptor signalling. Br. J. Pharmacol., 177(2), 449–463. https://doi.org/10.1111/bph.14879

Mark, K. S., Farquhar, B., Chisolm, M. S., Coleman-Cowger, V. H., & Terplan, M. (2015). Knowledge, attitudes, and practice of electronic cigarette use among pregnant women. J. Addict. Med, 9(4), 266–272. https://doi.org/10.1097/adm.0000000000000128

Matta, S. G., Balfour, D. J., Benowitz, N. L., Boyd, R. T., Buccafusco, J. J., Caggiula, A. R., Craig, C. R., Collins, A. C., Damaj, M. I., Donny, E. C., Gardiner, P. S., Grady, S. R., Heberlein, U., Leonard, S. S., Levin, E. D., Lukas, R. J., Markou, A., Marks, M. J., McCallum, S. E., Parameswaran, N., Perkins, K. A., Picciotto, M. R., Quik, M., Rose, J. E., Rothenfluh, A., Schafer, W. R., Stolerman, I. P., Tyndale, R. F., Wehner, J. M., & Zirger, J. M. (2007). Guidelines on nicotine dose selection for in vivo research. Psychopharmacology (Berl), 190(3), 269–319. https://doi.org/10.1007/s00213-006-0441-0

McCubbin, A., Fallin-Bennett, A., Barnett, J., & Ashford, K. (2017). Perceptions and use of electronic cigarettes in pregnancy. Health Educ. Res., 32(1), 22–32. https://doi.org/10.1093/her/cyw059

Metz, T. D., & Borgelt, L. M. (2018). Marijuana use in pregnancy and while breastfeeding. Obstet Gynecol, 132(5), 1198–1210. https://doi.org/10.1097/aog.0000000000002878

Montanari, C., Kelley, L. K., Kerr, T. M., Cole, M., & Gilpin, N. W. (2020). Nicotine e-cigarette vapor inhalation effects on nicotine & cotinine plasma levels and somatic withdrawal signs in adult male wistar rats. Psychopharmacology, 237(3), 613–625. https://doi.org/10.1007/s00213-019-05400-2

Nashed, M. G., Hardy, D. B., & Laviolette, S. R. (2020). Prenatal cannabinoid exposure: Emerging evidence of physiological and neuropsychiatric abnormalities. Front Psychiatry, 11, 1–46. https://doi.org/10.3389/fpsyt.2020.624275

Natale, B. V., Gustin, K. N., Lee, K., Holloway, A. C., Laviolette, S. R., Natale, D. R., & Hardy, D. B. (2020). ∆9-tetrahydrocannabinol exposure during rat pregnancy leads to symmetrical fetal growth restriction and labyrinth-specific vascular defects in the placenta. Sci. Rep., 10(1), 1–15. https://doi.org/10.1038/s41598-019-57318-6

Newsom, R., & Kelly, S. J. (2008). Perinatal delta-9-tetrahydrocannabinol exposure disrupts social and open field behavior in adult male rats. Neurotoxicol Teratol, 30(3), 213–219. https://doi.org/10.1016/j.ntt.2007.12.007

Nguyen, J. D., Aarde, S. M., Vandewater, S. A., Grant, Y., Stouffer, D. G., Parsons, L. H., Cole, M., & Taffe, M. A. (2016). Inhaled delivery of δ(9)-tetrahydrocannabinol (thc) to rats by e-cigarette vapor technology. Neuropharmacology, 109, 112–120. https://doi.org/10.1016/j.neuropharm.2016.05.021

Paul, S. E., Hatoum, A. S., Fine, J. D., Johnson, E. C., Hansen, I., Karcher, N. R., Moreau, A. L., Bondy, E., Qu, L., & Carter, E. B. (2019). Prenatal cannabis exposure and childhood outcomes: Results from the abcd study. Med-Archive, 78(1), 64–76. https://doi.org/10.1001/jamapsychiatry.2020.2902

Peters, D. A., & Tang, S. (1982). Sex-dependent biological changes following prenatal nicotine exposure in the rat. Pharmacol Biochem Behav., 17(5), 1077–1082. https://doi.org/10.1016/0091-3057(82)90497-x

Polli, F. S., & Kohlmeier, K. A. (2020). Prenatal nicotine exposure in rodents: Why are there so many variations in behavioral outcomes? N&TR, 22(10), 1694–1710. https://doi.org/10.1093/ntr/ntz196

Polli, F. S., Scharff, M. B., Ipsen, T. H., Aznar, S., Kohlmeier, K. A., & Andreasen, J. T. (2020). Prenatal nicotine exposure in mice induces sex-dependent anxiety-like behavior, cognitive deficits, hyperactivity, and changes in the expression of glutamate receptor associated-genes in the prefrontal cortex. Pharmacol Biochem Behav., 195, 1–11. https://doi.org/10.1016/j.pbb.2020.172951

Riedel, G., & Davies, S. (2005). Cannabinoid function in learning, memory and plasticity. Cannabinoids, 168(1), 445–477. https://doi.org/10.1007/3-540-26573-2_15

Römer, P., Mathes, B., Reinelt, T., Stoyanova, P., Petermann, F., & Zierul, C. (2020). Systematic review showed that low and moderate prenatal alcohol and nicotine exposure affected early child development. Acta Paediatr, 109(12), 2491–2501. https://doi.org/10.1111/apa.15453

Roncero, C., Valriberas-Herrero, I., Mezzatesta-Gava, M., Villegas, J. L., Aguilar, L., & Grau-López, L. (2020). Cannabis use during pregnancy and its relationship with fetal developmental outcomes and psychiatric disorders. A systematic review. Reprod. Health, 17(1), 1–9. https://doi.org/10.1186/s12978-020-0880-9

Scher, M. S., Richardson, G. A., Coble, P. A., Day, N. L., & Stoffer, D. S. (1988). The effects of prenatal alcohol and marijuana exposure: Disturbances in neonatal sleep cycling and arousal. Pediatr. Res., 24(1), 101–105. https://doi.org/10.1203/00006450-198807000-00023

Shacka, J., Fennell, O., & Robinson, S. (1997). Prenatal nicotine sex-dependently alters agonist-induced locomotion and stereotypy. Neurotoxicol Teratol, 19(6), 467–476. https://doi.org/10.1016/s0892-0362(97)00063-9

Slotkin, T. A., Epps, T. A., Stenger, M. L., Sawyer, K. J., & Seidler, F. J. (1999). Cholinergic receptors in heart and brainstem of rats exposed to nicotine during development: Implications for hypoxia tolerance and perinatal mortality. Brain Res. Dev. Brain Res., 113(1-2), 1–12. https://doi.org/10.1016/s0165-3806(98)00173-4

Smith, A. M., Mioduszewski, O., Hatchard, T., Byron-Alhassan, A., Fall, C., & Fried, P. A. (2016). Prenatal marijuana exposure impacts executive functioning into young adulthood: An fmri study. Neurotoxicol Teratol, 58, 53–59. https://doi.org/10.1016/j.ntt.2016.05.010

Sorenson, C. A., Raskin, L. A., & Suh, Y. (1991). The effects of prenatal nicotine on radial-arm maze performance in rats. Pharmacol Biochem Behav., 40(4), 991–993. https://doi.org/10.1016/0091-3057(91)90117-k

Stroud, L. R., Papandonatos, G. D., McCallum, M., Kehoe, T., Salisbury, A. L., & Huestis, M. A. (2018). Prenatal tobacco and marijuana co-use: Impact on newborn neurobehavior. Neurotoxicol Teratol, 70, 28–39. https://doi.org/10.1016/j.ntt.2018.09.003

Taghavi, T., Arger, C. A., Heil, S. H., Higgins, S. T., & Tyndale, R. F. (2018). Cigarette consumption and biomarkers of nicotine exposure during pregnancy and postpartum. Addiction, 113(11), 2087–2096. https://doi.org/10.1111/add.14367

Takahashi, K. A., & Linden, D. J. (2000). Cannabinoid receptor modulation of synapses received by cerebellar purkinje cells. J. Neurophysiol., 83(3), 1167–1180. https://doi.org/10.1152/jn.2000.83.3.1167

Thompson, R., Dejong, K., & Lo, J. (2019). Marijuana use in pregnancy. Obstet Gynecol Surv. 74(7), 415–428. https://doi.org/10.1097/ogx.0000000000000685

Tiesler, C. M. T., & Heinrich, J. (2014). Prenatal nicotine exposure and child behavioural problems. Eur Child Adolesc Psychiatry, 23(10), 913–929. https://doi.org/10.1007/s00787-014-0615-y

Tizabi, Y., Russell, L. T., Nespor, S. M., Perry, D. C., & Grunberg, N. E. (2000). Prenatal nicotine exposure. Pharmacol Biochem Behav., 66(3), 495–500. https://doi.org/10.1016/s0091-3057(00)00171-4

Turner, J. R. (2005). Nicotinic cholinergic receptors in the rat cerebellum: Multiple heteromeric subtypes. J. Neurosci, 25(40), 9258–9265. https://doi.org/10.1523/jneurosci.2112-05.2005

Vardaris, R. M., Weisz, D. J., Fazel, A., & Rawitch, A. B. (1976). Chronic administration of delta-9-tetrahydrocannabinol to pregnant rats: Studies of pup behavior and placental transfer. Pharmacol Biochem Behav, 4(3), 249–254. https://doi.org/10.1016/0091-3057(76)90236-7

Weimar, H. V., Wright, H. R., Warrick, C. R., Brown, A. M., Lugo, J. M., Freels, T. G., & McLaughlin, R. J. (2020). Long-term effects of maternal cannabis vapor exposure on emotional reactivity, social behavior, and behavioral flexibility in offspring. Neuropharmacology, 179, 1–38. https://doi.org/10.1016/j.neuropharm.2020.108288

Wickstrom, R. (2007). Effects of nicotine during pregnancy: Human and experimental evidence. Curr. Neuropharmacol, 5(3), 213–222. https://doi.org/10.2174/157015907781695955

Willford, J. A., Chandler, L. S., Goldschmidt, L., & Day, N. L. (2010). Effects of prenatal tobacco, alcohol and marijuana exposure on processing speed, visual–motor coordination, and interhemispheric transfer. Neurotoxicol. Teratol., 32(6), 580–588.

Young-Wolff, K. C., Sarovar, V., Tucker, L.-Y., Conway, A., Alexeeff, S., Weisner, C., Armstrong, M. A., & Goler, N. (2019). Self-reported daily, weekly, and monthly cannabis use among women before and during pregnancy. JAMA, 2(7), 1–10. https://doi.org/10.1001/jamanetworkopen.2019.6471

